# A novel *in vitro Caenorhabditis elegans* transcription system

**DOI:** 10.1101/2020.06.22.149609

**Authors:** Phillip Wibisono, Yiyong Liu, Jingru Sun

## Abstract

*Caenorhabditis elegans* is an excellent model organism for biological research, but its contributions to biochemical elucidation of eukaryotic transcription mechanisms have been limited. One of the biggest obstacles for biochemical studies of *C. elegans* is the high difficulty of preparing functionally active nuclear extract due to its thick surrounding cuticle. By employing Balch homogenization, we have achieved effective disruption of larval and adult worms and have obtained functionally active nuclear extract through subcellular fractionation. *In vitro* transcription reactions were successfully re-constituted using such nuclear extract. Furthermore, two non-radioactive detection methods, PCR and qRT-PCR, have been adapted into our system to qualitatively and quantitatively detect transcription, respectively. Using this system to assess how pathogen infection affects *C. elegans* transcription revealed that *Pseudomonas aeruginosa* infection increased transcription activity. Our *in vitro* system is useful for biochemically studying *C. elegans* transcription mechanisms and gene expression regulations. The effective preparation of functionally active nuclear extract in our system fills a technical gap in biochemical studies of *C. elegans* and will expand the usefulness of this model organism in addressing many biological questions beyond transcription.

## Background

*Caenorhabditis elegans* is a free-living, 1-millimeter-long, nematode worm found in soil and decaying organic matter. In 1963, Sydney Brenner proposed research into *C. elegans*, stating that “I would like to tame a small metazoan organism to study development directly” (1). Since then, *C. elegans* has been used as a model organism to address a wide range of biological questions such as those relating to development, metabolism, neurobiology, and aging. The nematode has many characteristics that make it an excellent model system, including, but not limited to, its rapid (3-day) life cycle, small size, ease of laboratory cultivation, genetic tractability, invariant lineage, effectiveness of RNA interference, and transparent body that allows for monitoring development or gene expression with single-cell resolution. Research involving the use of *C. elegans*, including Brenner’s work on organ development, was awarded with the Nobel Prize in 2002, 2006, and 2008.

Because of its genomic simplicity and physical characteristics, *C. elegans* offers a unique system to study transcription mechanisms and regulation. For example, mutations in pre-initiation complex genes have been recovered in genetic screens of *C. elegans* and have linked regulation that involves these factors to specific biological processes (2); the nematode is transparent throughout its entire life cycle making it an ideal system to use fluorescent protein reporter genes to monitor gene expression in live animals (3). Despite these advantages, however, the nematode’s contributions to biochemical elucidation of eukaryotic transcription mechanisms have been limited, whereas other model systems, such as *Saccharomyces cerevisiae, Drosophila melanogaster*, and cultured mammalian cells, were among the major contributors (2). In fact, very few biochemical studies of *C. elegans* transcription have been performed. One of the biggest obstacles for such studies is the high difficulty of obtaining functionally active nuclear extract due to the nematode’s thick surrounding cuticle; therefore, most analyses of transcription mechanisms in *C. elegans* have employed intact embryos or whole animals (2). The nematode cuticle is a highly structured extra-cellular matrix comprising predominantly cross-linked collagens, additional insoluble proteins termed cuticlins, and associated glycoproteins and lipids (4). It is synthesized five times during development, once in the embryo and subsequently at the end of each larval stage prior to molting, making *C. elegans* at all stages resistant to buffer extraction or mechanic forces (4). A *C. elegans in vitro* transcription system was once developed by Lichtsteiner and Tjian in the 1990s (5, 6), but it has not become widely used, most likely because the transcription reactions were re-constituted with nuclear extract from embryos, not from larval or adult worms, and the method of Dounce homogenization used to prepare the extract could lead to protein instability (7). Besides Dounce homogenization, several other techniques have also been described to break worms, including pressure cycling technology (PCT), bead beating, grinding after flash cooling, sonication, and Balch homogenization (7–11). Most of these methods have their own advantages and disadvantages and there are no reports of transcription reactions being re-constituted following worm disruption using these approaches. A *C. elegans* transcription system with effective preparation of functionally active nuclear extract from larval or adult worms has yet to be established.

Eukaryotic transcription is a complex biochemical process catalyzed by three nuclear RNA polymerases that synthesize different types of RNA: RNA polymerase I (Pol I) catalyzes the transcription of all rRNA genes except 5S; Pol II synthesizes mRNAs and many non-coding RNAs, including small nuclear RNAs; and Pol III makes tRNAs and other small non-coding RNAs, including 5S rRNA and U6 small nuclear RNA (12). While RNA Pol I and Pol III have been studied very little in *C. elegans*, the Pol II-mediated mRNA transcription appears to be conserved in the nematode (2). As in other eukaryotes, mRNA transcription in *C. elegans* includes three phases: initiation, elongation, and termination. The first step is assembly of a pre-initiation complex on promoter DNA, followed by DNA opening and synthesis of a short initial RNA oligomer. Pol II then uses the DNA template to extend the growing RNA chain in a processive manner. Finally, DNA and RNA are released during termination and Pol II can then be recycled to re-initiate transcription. These basic mRNA transcription mechanisms are believed to be conserved in *C. elegans* largely because most of the core transcription factors are conserved in *C. elegans* at the DNA sequence level, and because *in vivo* studies of the nematode’s transcription machinery components have generally yielded results consistent with mechanistic functions that were defined by *in vitro* studies of other systems (2). Establishing an *in vitro C. elegans* transcription system is necessary to biochemically identify similarities and differences in transcription between *C. elegans* and other eukaryotes, and, more importantly, to further elucidate transcription mechanisms and gene expression regulations.

In the current study, we have developed an *in vitro C. elegans* transcription system with effective disruption of larval or adult worms and preparation of functionally active nuclear extract. Traditionally, the detection of transcription activity has relied on radioactive labeling of the newly synthesized RNA and visualization of incorporated radioactivity by autoradiography (13). However, radioisotope labels have many drawbacks including perceived health concerns, regulatory requirements, disposal problems, and short shelf life (14). While technological advancements have allowed the use of non-radioactive methods for RNA detection, such as quantitative reverse transcription PCR (qRT-PCR) (15) or the use of fluorescent nucleotides to label newly synthesized RNA (16), these new methods have been mainly developed to investigate transcription in mammalian cell models. Here, we have adapted two non-radioactive methods, namely PCR and qRT-PCR, for qualitatively and quantitatively detecting RNA in our *in vitro* system, respectively. By coupling *in vitro* transcription reactions using our *C. elegans* nuclear extract preparations with these non-radioactive RNA detection approaches, we have examined how pathogen infection affects *C. elegans* transcription and revealed that infection with *Pseudomonas aeruginosa* strain PA14, a human opportunistic pathogen, increased the nematode’s transcription activity. Overall, we have shown that our *in vitro* system can be useful for biochemical studies of *C. elegans* transcription. Our approach for effective disruption of larval or adult worms and preparation of functionally active nuclear extract could also expand the usefulness of the *C. elegans* model to biochemically address other biological questions.

## Results

### Preparation of *C. elegans* nuclear extract

Although subcellular fractionation is widely used to prepare nuclear extract from cultured cells or animal tissues, subcellular fractionation of *C. elegans* has been challenging because of the worms’ thick surrounding cuticle. Several techniques have been described to break worms, including pressure cycling technology (PCT), bead beating, grinding after flash cooling, sonication, Dounce homogenization, and Balch homogenization (7–11); however, most of these methods have known advantages and disadvantages. For example, the PCT can achieve complete disruption of worms through high pressure, but it requires expensive equipment and mixing of animals with an SiC abrasive; bead beating is cheap and fast, but it results in highly variable protein recoveries as well as protein aggregation and denaturation; grinding after flash cooling is rapid, but it needs a large amount of starting materials due to significant sample loss; sonication and Dounce homogenization are effective but could lead to protein instability. In contrast, Balch homogenization, which uses a proprietary device Balch homogenizer (Fig. 1A), has been shown by Bhaskaran *et al.* (7) to effectively disrupt worms and yield functional protein extracts. This method also generates amounts of soluble proteins that are comparable to sonication or Dounce homogenization, while also keeping proteins, nucleic acids, mitochondria, and polysomes intact (7). Because of these advantages and the device’s affordable price (~$2,000 each), we decided to adapt Balch homogenization for nuclear extraction in our *C. elegans* transcription system.

**Fig. 1.**
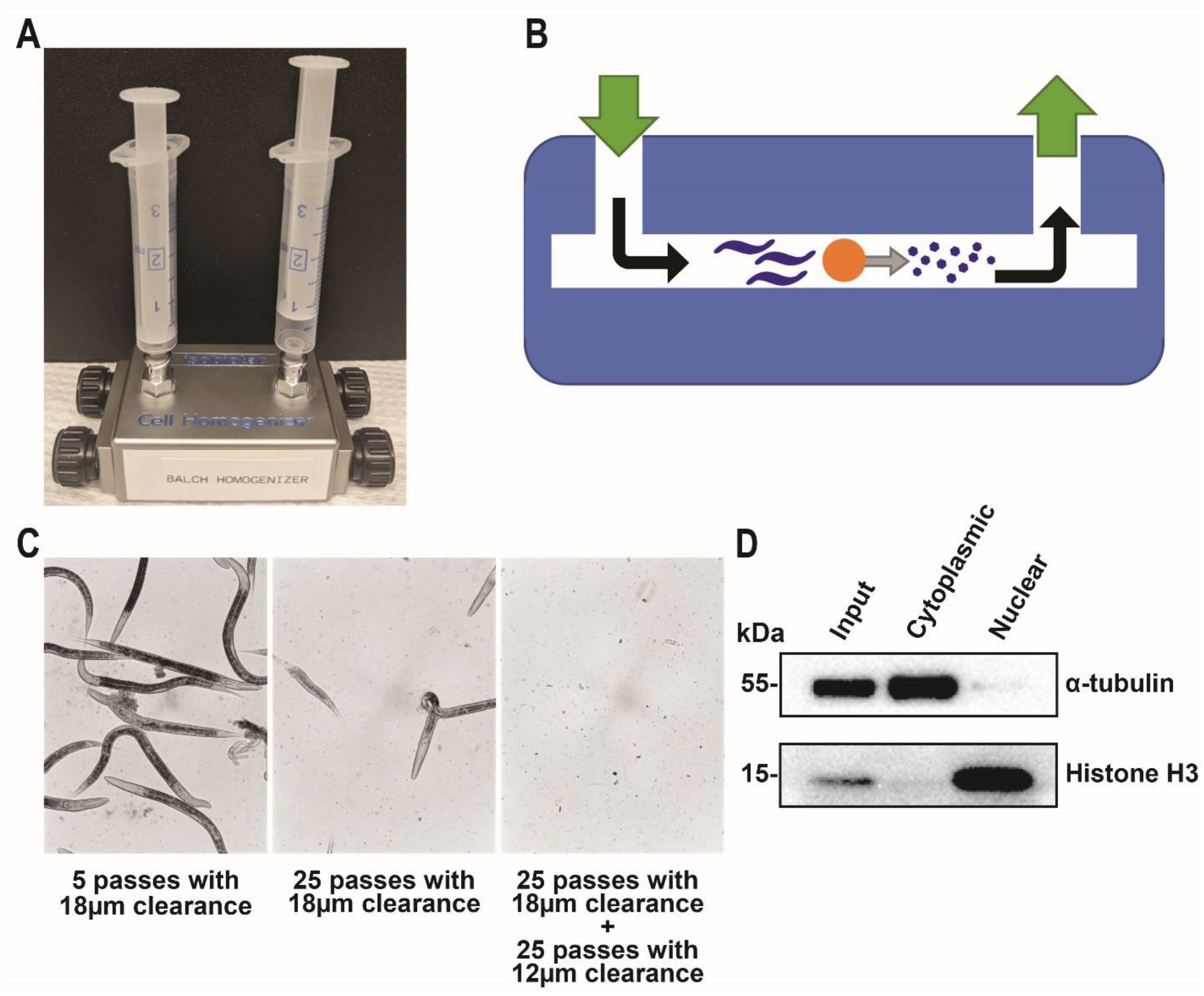
Preparation of *C. elegans* nuclear extract. **(A)** A photo of Balch homogenizer. **(B)** A schematic of Balch homogenizer. The chamber of Balch homogenizer was loaded with a ball bearing and worms suspended in complete hypotonic buffer. Worms were pushed through the gap between the chamber wall and the ball bearing. This movement was repeated to slowly break open the worms and produce a homogenate. **(C)** Worm lysates after different combinations of syringe pass numbers and ball-bearing sizes. **(D)** Western blot of fractionated homogenate. α-tubulin and histone H3 were probed as markers for the cytoplasmic fraction and nuclear extract, respectively. Input, 5 μg of homogenate before fractionation; cytoplasmic, 5 μg of the cytoplasmic fraction; nuclear, 5 μg of the nuclear extract.

A Balch homogenizer consists of a hollow metal chamber into which a ball bearing of defined diameter is inserted (a schematic is depicted in Fig. 1B). The gap or clearance between the chamber wall and the ball bearing is adjustable by using ball bearings of different diameters. Worms in a buffer can be repeatedly pushed through the gap back and forth from the two syringes connected to the two ends of the chamber to generate homogenized lysate (homogenate). The homogenizer can also be placed on ice during operation to keep the sample from overheating. We tested different combinations of syringe pass numbers and ball-bearing sizes for optimal homogenization and observed that 25 passes with an 18μm-clearance ball bearing followed by 25 passes with a 12μm-clearance ball bearing resulted in complete disruption of L4 larvae or young adult worms (Fig. 1C). This procedure was then used for all worm homogenization in the studies described below. After homogenization, the cytoplasmic fraction was separated from the nuclei by centrifugation, and the nuclei were lysed in a hypotonic buffer to obtain nuclear extract. We subsequently checked the effectiveness of our subcellular fractionation through Western blot analyses of the cytoplasmic fraction and nuclear extract using α-tubulin and histone H3 as marker proteins, respectively (17). Western results showed that α-tubulin mainly appeared in the cytoplasmic fraction and not in the nuclear fraction, whereas histone H3 was exclusively detected in the nuclear extract (Fig. 1D), indicating that proper subcellular fractionation was achieved.

### *In vitro* detection of *C. elegans* transcription by PCR

After succeeding in preparation of *C. elegans* nuclear extract, we proceeded to re-constitute transcription reactions *in vitro* using the extract. Traditional *in vitro* transcription assays rely on radioactive labeling of the newly synthesized RNA and detection of incorporated radioactivity by autoradiography (13). However, as technology has advanced, non-radioactive detection methods have been developed. Voss *et al.* reported a simple qualitative PCR detection of RNA transcripts (15), so we decided to adapt this method to detect transcription of *C. elegans* nuclear extract. Fig. 2A depicts the scheme of our experimental procedure. Briefly, a linear DNA containing the CMV promoter was used as the template (DNA sequence is listed in Table 1); nuclear extract was added to the DNA template to synthesize RNA, followed by RNA purification and reverse transcription; the resulting cDNA was amplified by PCR with the primer pair *HNqPCRrevl* and *HNqPCRfrwl* (Table 1) to produce a 132-bp DNA; the amplified DNA was then run on agarose gel, followed by staining and imaging. Because HeLa nuclear extract has been proven to work efficiently in *in vitro* transcription assays (18), we first set up transcription reactions using the HeLa Scribe Nuclear Extract *in vitro* Transcription System (Promega, catalog # E3110); however, instead of using radioactive nucleotides and autoradiography detection, we followed the above-described PCR protocol. As shown in Fig. 2B, a DNA fragment of expected size (132 bp) was produced, confirming that the PCR method can amplify and detect the RNA transcript. Substitution of HeLa nuclear extract with *C. elegans* nuclear extract in the transcription reactions generated a DNA product of the same size (Fig. 2B), indicating that the *C. elegans* nuclear extract was capable of re-constituting transcription and that the nematode can use the CMV promoter to transcribe DNA. We then attempted to adapt this PCR method to quantitatively measure the transcription activity of *C. elegans* nuclear extract by quantifying DNA in gel images using ImageJ software. Although the quantification results were consistent between replicates in the same experiments, large variations were observed between experiments (data not shown), which indicates that this PCR method is not a good quantitative approach, possibly due to the multiple steps involved in the quantification (*i.e.*, conversion of RNA into cDNA, amplification of cDNA by PCR, gel electrophoresis, and densitometric measurement using ImageJ). Taken together, we have successfully adapted a PCR method to qualitatively detect *C. elegans* transcription *in vitro*.

**Fig. 2.**
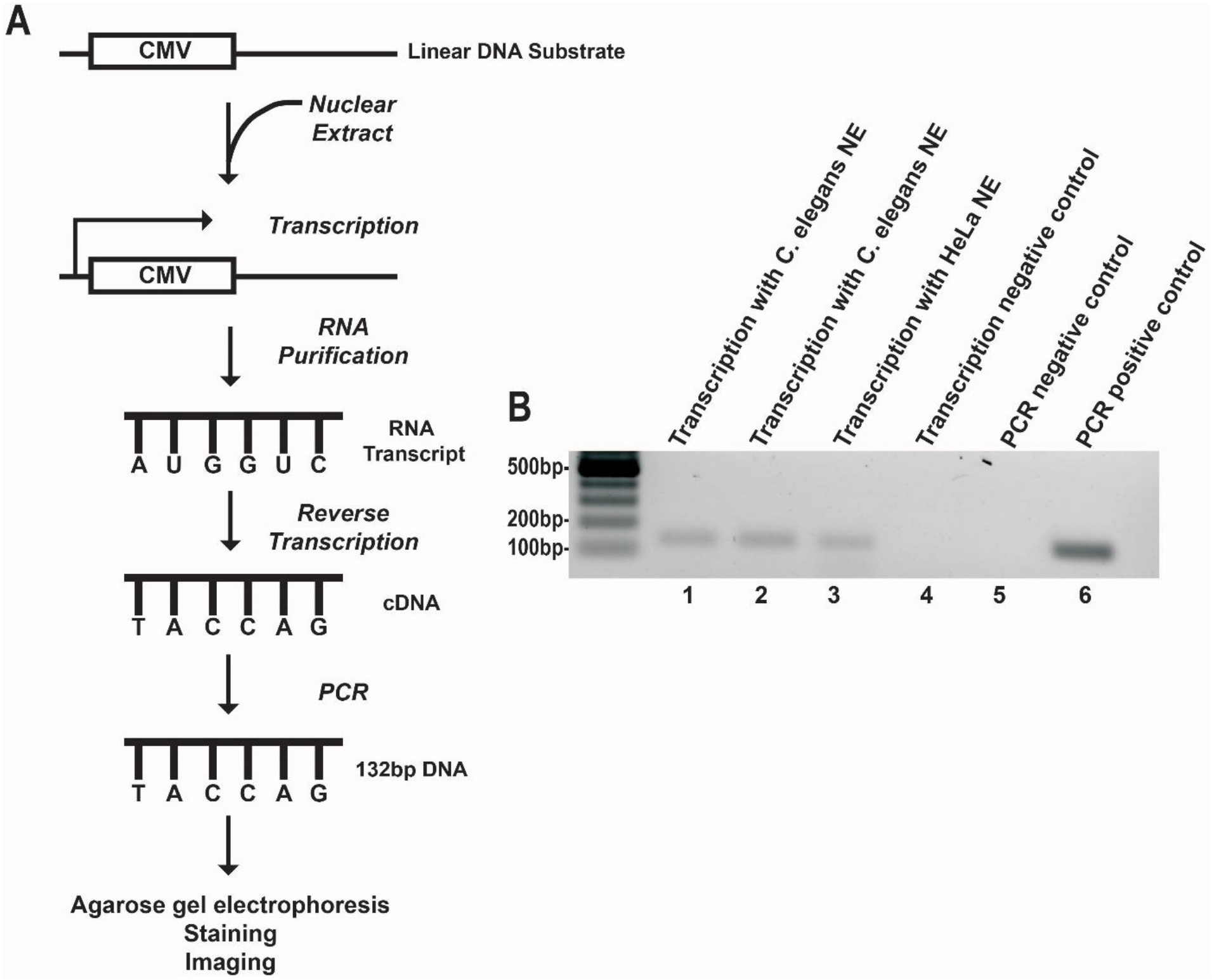
Detection of *C. elegans* transcription by PCR. **(A)** Scheme of PCR detection of *C. elegans* transcription. A linear DNA containing the CMV promoter was used as the template; nuclear extract was added into the reaction to synthesize RNA, followed by RNA purification and reverse transcription; the resulting cDNA was amplified by PCR; the DNA product was then run on agarose gel, stained with SYBR Safe Stain, and imaged with a CCD camera. **(B)** A gel image of PCR detection. Lanes 1 and 2, transcription with *C. elegans* nuclear extract; lane 3, transcription with HeLa nuclear extract; lane 4, transcription negative control (without any nuclear extract); lane 5, PCR negative control (without any nucleotides); lane 6, PCR positive control.

**Table 1:**
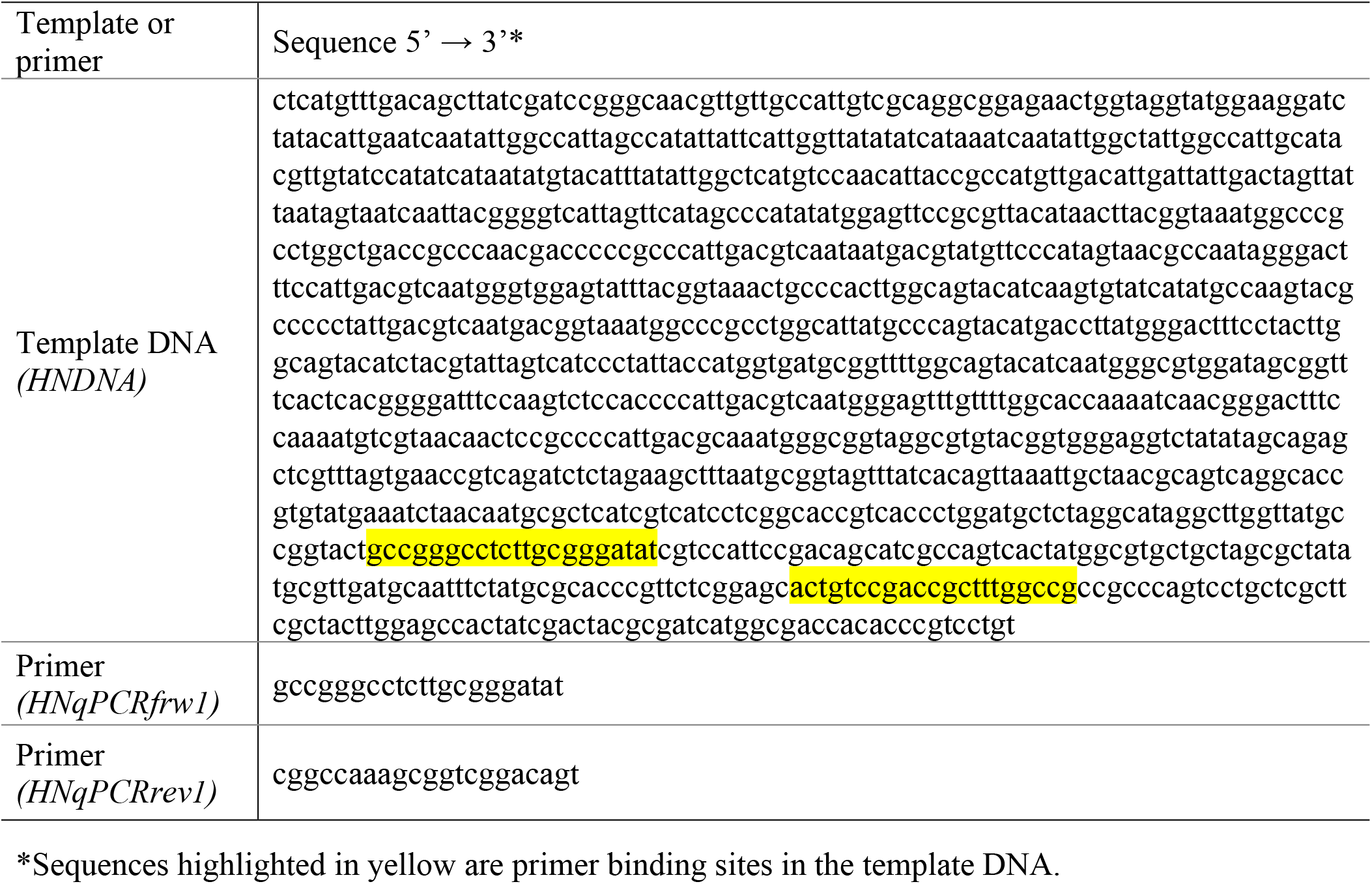
Sequences of template DNA and primers for PCR and qRT-PCR

### *In vitro* quantification of *C. elegans* transcription activity by qRT-PCR

We next sought to quantify *C. elegans* transcription activity using qRT-PCR, a more direct approach than the above-described PCR method because it amplifies DNA and simultaneously measures DNA amounts using a fluorescent reporter. To this end, we followed the same experimental procedure as depicted in Fig. 2A to generate cDNA and then amplified cDNA by qRT-PCR with SYBR green detection. To assess the quantifiability of this method, we first conducted transcription reactions using various amounts of HeLa nuclear extract followed by qRT-PCR. Because in qRT-PCR the intensity of fluorescent signal is directly proportional to the number of amplified DNA molecules (19), a standard curve of the template DNA was included in the assays and used for RNA copy number calculations with the assumption that every RNA molecule was reverse-transcribed into a DNA molecule (Fig. 3A). Conversion of fluorescent signals to RNA copy numbers and fitting of the latter with the Michaelis-Menten model and the non-linear least-squares method, as described previously (20), yielded a coefficient of determination (*r^2^*) of 0.9773 (Fig. 3B), indicating that the enzymatic reactions in transcription follows the Michaelis-Menten model and that the qRT-PCR method used allows for quantitative measurement of transcription activity. Substitution of HeLa nuclear extract with 5 μg of *C. elegans* nuclear extract in the transcription reactions produced 102,589 RNA molecules on average, which was 7-fold of what equivalent HeLa nuclear extract produced (Fig. 3B). Therefore, *C. elegans* nuclear extract was 6-fold more active in generating RNA than HeLa nuclear extract under our *in vitro* transcription conditions. This large difference in transcription between the two nuclear extracts could be due to several reasons. For example, the HeLa nuclear extract was from a commercial source (Promega, catalog # E3110) with an unknown preparation date, whereas our *C. elegans* nuclear extract was freshly made. The assay temperature was 30°C, which is not optimal for most mammalian enzymes (37°C is optimal for most mammalian enzymes (21)) but worked well for *C. elegans* enzymes.

**Fig. 3.**
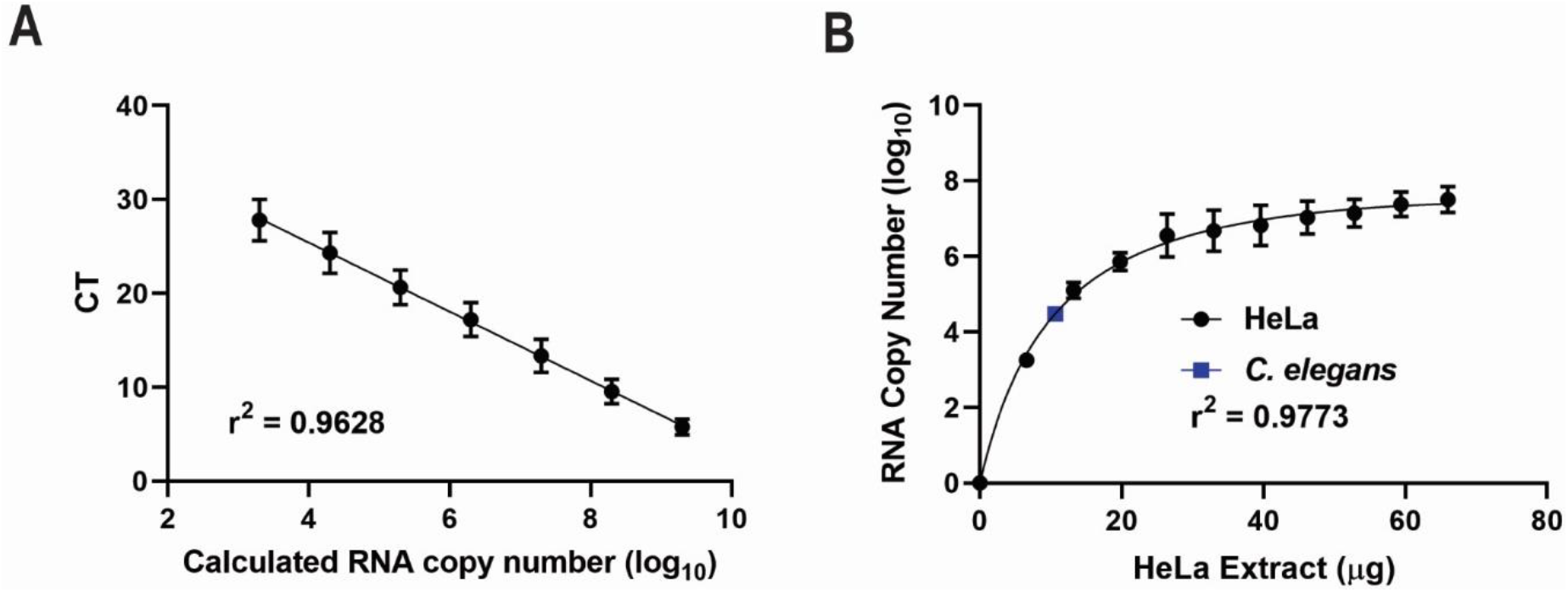
Quantitative analysis of RNA transcription by qRT-PCR. *In vitro* transcription was performed, and the resulting RNA was reverse transcribed into cDNA, followed by qRT-PCR quantification of the cDNA. **(A)** The template DNA was serially diluted 10-fold for seven times and subjected to qRT-PCR. The cycle thresholds (CTs) were plotted against the expected copy numbers of the template DNA, showing a linear relationship (Y = −3.677x + 40.12, *r^2^* = 0.9628). This standard curve was then used for the calculation of RNA copy number generated from transcription reactions. The graph represents the combined results of three independent experiments. Error bars represent standard deviation. **(B)** The RNA copy numbers generated from transcription were plotted against the amounts of HeLa nuclear extract used in the reactions, followed by fitting with the Michaelis-Menten model and the non-linear least-squares method in Excel spreadsheets (Y = 9.568*X/(11.34+X) - 0.01128*X - 0.02921, *r^2^* = 0.9773). Substitution of HeLa nuclear extract with 5 μg of *C. elegans* nuclear extract in the transcription reaction produced 102,589 RNA molecules on average. The graph represents the combined results of three independent experiments. Error bars represent standard deviation.

### Comparison of *C. elegans* transcription activity with or without *P. aeruginosa* infection

Next we applied the above-described transcription system to assess how pathogen infection affects *C. elegans* transcription activity. Upon infection, host organisms launch stress responses to fight invading microbes, in part, by altering gene expression (22). While many research efforts have been focused on investigating activation and/or inactivation of specific genes or signaling pathways in response to infection, how general RNA pol II transcription activity changes is not well understood. Our *in vitro C. elegans* transcription system offers a suitable approach to address this question at the whole-organism level. To this end, we chose the human opportunistic pathogen *P. aeruginosa* strain PA14 for worm infection, because the *P. aeruginosa-C. elegans* infection model is well-established for bacterial pathogenesis research (23–25). To infect worms, young-adult worms fed on standard worm food *Escherichia coli* strain OP50 were transferred to a lawn of *P. aeruginosa* and incubated at 25°C for 1 hour. Uninfected control worms stayed on *E. coli* at 25°C for 1 hour. These worms were then collected, nuclear extract was prepared using Balch homogenization, and transcription assays were conducted with qRT-PCR quantification, as described above. Results showed that on average, 1 μg of nuclear extract from uninfected worms transcribed 6,072 RNA molecules, while infected nuclear extract transcribed 9,837 RNA molecules (Fig. 4), a 0.62-fold increase upon infection (Fig. 4). This result demonstrated that *P. aeruginosa* infection increase overall transcription activity, supporting the view that *C. elegans* upregulates its transcription activity to meet increased demand for protein production in order to fight infection (26).

**Fig. 4.**
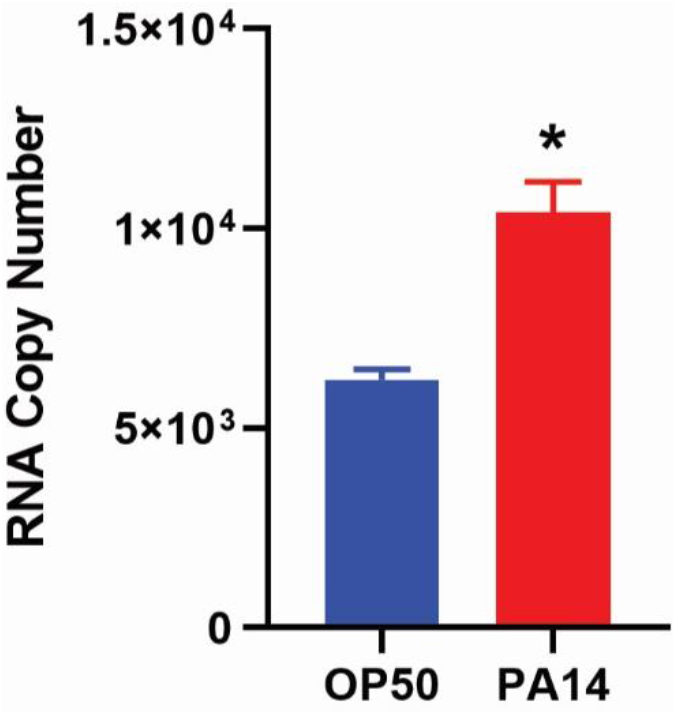
Comparison of *C. elegans* transcription activity with or without *P. aeruginosa* infection. Synchronized young-adult worms were exposed to *P. aeruginosa* PA14 or *E. Coli* OP50 as a control at 25°C for 1 hour. The worms were then collected, followed by nuclear extract preparation and transcription assays with qRT-PCR quantification. On average, 1 μg of nuclear extract from uninfected worms transcribed 6,072 RNA molecules, while infected nuclear extract transcribed 9,837 RNA molecules, a 0.62-fold increase upon infection. The graph represents the combined results of three independent experiments. Error bars represent standard deviation. Asterisk (*) denotes a significant difference (*P* < 0.05) between infected worms and uninfected control worms, as analyzed by a two-sample *t* test for independent samples.

## Discussion

Even though *C. elegans* has been used as a model system to address a wide range of biological questions, the nematode’s contributions to biochemical elucidation of eukaryotic transcription mechanisms have been limited. One of the biggest hurdles in *in vitro* studies of *C. elegans* transcription is the high difficulty of obtaining functionally active nuclear extract due to the nematode’s thick surrounding cuticle. By adapting the method of Balch homogenization (7), we have achieved effective worm disruption and optimal subcellular fractionation, resulting in a nuclear exact that is functionally active in inducing transcription appropriately. Two non-radioactive detection methods, PCR and qRT-PCR, have also been adapted into our transcription system. The PCR method can qualitatively detect *C. elegans* transcription *in vitro*, and the qRT-PCR method can quantitatively measure transcription activity. Applying this system to assess how pathogen infection affects *C. elegans* transcription revealed that *P. aeruginosa* infection increased its transcription activity. Therefore, our *in vitro* system can be a useful tool for biochemically studying transcription mechanisms and gene expression regulation in *C. elegans*, which will facilitate our understanding of transcription in higher organisms due to the conservation of eukaryotic transcription.

As in other eukaryotes, the RNA Pol II-mediated transcription in *C. elegans* depends on the binding of transcription factors to specific gene *cis*-acting sequences (3). These *cis*-regulatory sequences are usually clustered into discrete functional modules including the core promoter, extended proximal and downstream promoter regions, positive and negative enhancers, and insulators. Pol II acts in concert with TATA Binding Protein (TBP) and TBP-Associated Factors (TAFs) at the core promoter to initiate transcription. The core promoter in *C. elegans* typically includes five elements: an Sp1 like site (CNCCGCCC), T-blocks that correlate with nucleosome eviction and gene expression levels (TTTT[N/T]), a TATA box (GTATA[TA][TA]AG), a trans-splicing site (TTnCAG), and a Kozak site that includes the translation initiation codon ([CA]AA[CA]ATG) (27). We employed a simple DNA substrate in our *in vitro* system that contains the CMV promoter and other minimal elements required to induce transcription using nuclear extract from mammalian cells (15). Its core promoter contains all five elements (or similar sequences) typical for *C. elegans* and was able to induce transcription appropriately with nematode nuclear extract. These results suggest that, like in mammals, the CMV promoter can regulate gene expression in *C. elegans*. This *in vitro* system will be useful for biochemically studying *C. elegans* transcription machinery components. For example, because a custom DNA template is used in this system, the role of any *cis*-acting sequence can be investigated by including or excluding the specific sequence in the template. Similarly, transcription factor functions can be studied by depleting or complementing the nuclear extract with specific factors. This system should be especially suitable for examining the interactions of transcription factors and *cis*-acting sequences because the interacting complexes can be pulled down by using either specific antibodies against the proteins or streptavidin beads against a biotinylated DNA template. Many mechanistic questions regarding *C. elegans* transcription that could not be answered previously can now be biochemically interrogated using our system. However, one drawback of this system is that spatial regulation of transcription, such as cell- or tissue-specific gene expression, cannot be investigated because the nuclear extract is prepared from whole animals.

The usefulness of our transcription system in solving biological problems has been demonstrated by the comparison of *C. elegans* transcription activity with or without *P. aeruginosa* infection, which revealed that infection increases the nematode’s transcription activity. Although this does not necessarily reflect cell- or tissue-specific transcriptional response to pathogen infection, it provides a global view of infection-induced transcriptional changes at the whole-organism level and biochemically supports the notion that *C. elegans* upregulates its transcription activity to meet increased demand for protein production to fight infection (26). By the same token, this system can be used to investigate global transcriptional responses to other environmental or internal assaults such as chemical toxins, ultraviolet radiation, genetic mutations, treatments of diseases, or aging. Therefore, our *in vitro* transcription system not only fills a technical gap in biochemical studies of *C. elegans*, but also expand the usefulness of this powerful model organism in addressing many biological questions.

## Conclusions

In this study, we have developed an *in vitro C. elegans* transcription system that re-constitutes transcription reactions using nuclear extract of larval or adult worms, and can both qualitatively and quantitatively detect transcription activity using non-radioactive approaches. This *in vitro* system employs Balch homogenization to effectively disrupt worms followed by subcellular fractionation, resulting in a nuclear extract that is functionally active in inducing transcription appropriately. We have adapted two non-radioactive detection methods, PCR and qRT-PCR, to qualitatively and quantitatively, respectively, detect *C. elegans* transcription *in vitro*. Overall, this system will be useful for biochemically studying *C. elegans* transcription mechanisms and gene expression regulations. More specifically, it can be used to study *C. elegans* transcription machinery components, such as transcription factors, *cis*-acting sequences, or interactions between them; it can also be used to examine gene expression regulations under specific environmental or internal conditions. Knowledge gained from these studies will also facilitate our understanding of transcription in higher organisms due to the conservation of eukaryotic transcription. Furthermore, the ability to effectively prepare functionally active nuclear extract for use in our *in vitro* system fills a technical gap in biochemical studies of *C. elegans* and will expand the usefulness of this powerful model organism in addressing many biological questions beyond transcription.

## Methods

### Bacterial and *C. elegans* strains

*E. coli* OP50 and *P. aeruginosa* PA14 were used in this study. These bacteria were grown in Luria-Bertani (LB) broth at 37°C. Wild-type *C. elegans* Bristol N2 was used and cultured under standard conditions (28).

### Worm disruption and subcellular fractionation

Gravid adult animals were lysed using a solution of sodium hydroxide and bleach (ratio 5:2), and eggs were synchronized for 22 hours in S-basal liquid medium at room temperature. Synchronized L1 larval animals were transferred onto NGM plates seeded with *E. coli* OP50 and grown at 20°C for 48 hours until the animals reached L4 larval stage or for 72 hours until young adult stage. The animals were then collected, washed with M9 buffer, and used for nuclear extract preparation directly or for pathogen infection followed by nuclear extract preparation. For pathogen infection, the collected worms were transferred to NGM plates containing *E. coli* OP50 or *P. aeruginosa* PA14 at 25°C and incubated for 1 hour. Following this incubation, the animals were collected and washed with M9 buffer three times. The animals were then washed again in 3ml cold hypotonic buffer (15mM HEPES KOH, pH 7.6, 10mM KCl, 5 mM MgCl2, 0.1mM EDTA, and 350mM Sucrose) and centrifuged at 1425 x g for 3 minutes at room temperature. The supernatant was discarded, and the animal pellet was resuspended in 1ml of cold hypotonic buffer with 2x Protease inhibitor (ThermoFisher, catalog # 78430). The animals were transferred to a Balch homogenizer and grounded using an 18μm clearance ball bearing for 25 passes, followed by another 25 passes using a 12μm ball bearing. The final animal homogenate was transferred to a 1.5ml microtube and centrifuged at 500 x g for 5 minutes at 4°C. The supernatant was then transferred to a new 1.5ml microtube, and 40μl of supernatant was aliquoted to a microtube labeled “input”. The remaining supernatant was centrifuged at 4000 x g for 5 minutes at 4°C to pellet the nuclei. The supernatant was transferred to a new microtube and centrifuged at 17,000 x g for 30 minutes at 4°C, and the supernatant was collected as the “cytoplasmic fraction”. The nuclei pellet was washed with 500μl of complete hypotonic buffer and centrifuged at 4000 x g for 5 minutes at 4°C. The supernatant was discarded, and the nuclei pellet was resuspended in 500μl of complete hypotonic buffer and transferred to a new microtube. The resuspended nuclei was then centrifuged at 4000 x g for 5 minutes at 4°C. After discarding the supernatant, the nuclei pellet was resuspended in 40μl cold hypertonic buffer (15 mM HEPES KOH, pH 7.6, 400 mM KCl, 5mM MgCl2, 0.1mM EDTA, 0.1% Tween20, 10% Glycerol, 2x Protease inhibitor) and the suspension was transferred to a new microtube as the “nuclear fraction”. Fractions were quantified using Invitrogen Qubit protein assay (ThermoFisher, catalog # Q33221) before being snap frozen. All samples were stored at −80°C until use in the *in vitro* transcriptional assay.

### Western blot

Western blotting was performed as previously described (29). Briefly, equal mass aliquots of isolated cytoplasmic, nuclear, and input fractions were diluted with H2O to 7.5μl, mixed with 2.5μl of NuPAGE LDS buffer (Invitrogen, catalog # NP0007), and heated to 70°C for 10 minutes for denaturation. The isolates were separated on a NuPAGE 4-12% Bis-tris gel and transferred to a PVDF membrane. The primary antibodies used were mouse anti-tubulin alpha antibody (AA43) developed by Walsh, C (obtained from the Developmental Studies Hybridoma Bank at the University of Iowa, Department of Biology) and rabbit anti-histone H3 antibody (Novus Biologicals, catalog # NB500-171). The secondary antibodies used were goat anti-mouse IgG (H+L) antibody conjugated to HRP (Invitrogen, catalog # 31430) and goat anti-rabbit IgG (H+L) antibody conjugated to HRP (Promega, catalog # W4018). Immunoblots were imaged using iBright 1500 (ThermoFisher, catalog # A44241).

### *In vitro* transcription assay and RNA purification

Transcription assays were set up using the HeLa Scribe Nuclear Extract *in vitro* Transcription system (Promega, catalog # E3110) following the manufacture’s protocol with modifications. A master mix was prepared to include 1.5μl of 50mM MgCl2, 1.0μl of rNTPs (10mM of each), 4μl of linear HNDNA (25ng/μl), and 7.5μl of RNase-free H2O. Assay tubes were filled with 5μg of nuclear extract and transcription buffer to a total volume of 11μl. Fourteen μl of master mix was transferred to each assay tube, and the reactions were incubated at 30°C for 30 min. Two control reactions were performed: one containing 8 units of HeLa nuclear extract provided by Promega as a positive control and the other containing no nuclear protein as a negative control. Reactions were halted with the addition of 400μl of RLT buffer from the RNeasy Micro kit (Qiagen, catalog # 74004), snap frozen, and stored at −80°C until RNA cleanup. RNA cleanup was done using the RNeasy Micro kit following the manufacture’s recommendations, including on-column DNase treatment but excluding the addition of carrier RNA. The purified RNA was eluted from the column using 17μl of RNase-free H2O. Eluted RNA was again treated with DNase using the Baseline-ZERO DNase kit (Lucigen, catalog # DB0715K) following the manufacturer’s instructions. All RNA samples were stored at −80°C until reverse transcription.

### PCR amplification of transcription products

Reverse transcription reactions containing 2μl of purified RNA and 2μl of 10mM *HNqPCRrev1* primer in a final volume of 20μl were performed using the Qiagen Sensiscript RT kit (Qiagen, catalog # 205211) per the manufacturer’s instructions. After the reaction was completed, 1μl of the reverse transcription mix was transferred to a PCR tube and used for PCR. The standard PCR reaction was performed with the *HNqPCRrev1* and *HNqPCRfrw1* primers using the Failsafe PCR system with the Premix A (Lucigen, catalog # F599100). Positive and negative PCR control reactions with or without 50ng of HNDNA, respectively, were also performed. After amplification, the PCR products were analyzed by gel electrophoresis on a 2.0% agarose TAE gel, followed by staining with SYBR safe DNA gel stain (Invitrogen, catalog # S33102), imaging with iBright 1500, and quantification with software ImageJ.

### qRT-PCR

Quantification by qRT-PCR was done on the StepOnePlus 96-well real-time PCR system (Applied Biosystems, catalog # 4376600) using the PowerUp SYBR green qPCR kit (Applied Biosystems, catalog # A25918). Two microliters of reverse transcription product were amplified using 500nM each of the *HNqPCRrev1* and *HNqPCRfrw1* primers in a 10μl reaction. A seven-point standard curve of the linear HNDNA was included in every qRT-PCR experiment and used for RNA copy number calculations with the assumption that every RNA molecule was reverse-transcribed into a DNA molecule. Titration of HeLa nuclear extract standards was performed using the *in vitro* transcription assay followed by qRT-PCR quantification. The DNA products were converted to RNA copy numbers using the standard curve of the linear HNDNA and then fitted with the Michaelis-Menten model and the non-linear least-squares method in Excel spreadsheets, as described previously (20). The resulting fitting equation was used for calculating *C. elegans* nuclear extract transcription activity.

## List of abbreviations

PCR: polymerase chain reaction
qRT-PCR: quantitative reverse transcription polymerase chain reaction
NGM plates: nematode growth media plates
CMV: Cytomegalovirus
HNDNA: HeLa Nuclear DNA Template

## Declarations

### Authors’ contributions

PW designed and performed experiments, analyzed data, and was a major contributor in writing the manuscript. YL designed experiments, analyzed data, and was a major contributor in writing the manuscript. JS designed experiments, analyzed data, and was a major contributor in writing the manuscript. All authors read and approved the final manuscript.

